# Quantification of bioagressors induced yield gap for grain crops in France

**DOI:** 10.1101/641563

**Authors:** Nathan G.F. Devaud, Corentin M. Barbu

## Abstract

Analyzing yield losses of crops is instrumental in sustaining high productivity. Here, we develop statistical modelling of yield losses by French department based on weather, soil and epidemiological data for 13 diseases and 12 insect pests of grain crops over 8 years. Those environmental factors explain up to 90% of yield variations (wheat). Weather and soil quality are first order determinants as manifested both by direct assessment and by strongly correlated yields of even taxonomically very different crops. Bioagressors are important second order determinants: losses of 5 qx/ha (~7%) on wheat and 2 qx/ha (~15%) on winter oilseed rape. Across models based on conceptually different achievable yields, only *Septoria tritici* and *Sitobion avenae* on wheat, and *Ceutorhynchus picitarsis* and *Ceutorhynchus assimilis* on winter oilseed rape are consistently and significantly detrimental to yield. Those bioagressors seem not fully controlled which is compatible with empirical observations in the non academical litterature.

**Highlights:**

- Grain crop yields of similar growing seasons are highly correlated
- Environmental determinants explain up to 90% of yield variations
- Bioagressors induced yield variations of 5 to 20%
- Specific pests and diseases with less than perfect control are identified

## 1. Introduction

Determining average crop losses would allow setting priorities for research to maintain high productiv^it^y [1]. Monitoring losses is also important as public scrutiny on pesticides leads to more and more control products to be banned [2]. Field crops represent more than XXX% of the cultivated areas in France, account for a substantial share of French exports and represent more than half of pesticide use (http://agreste.agriculture.gouv.fr/page-d-accueil/article/agreste-donnees-en-ligne)Faire *une réf en bonne et due forme* making both a strategic ressource and a important lever to reduce pesticide use. Despite their importance, the average impact of the grain crop bioagressors on the yields is difficult to assess.

Studies over a few years may not assess correctly the harmfulness of bioagressors because abundances may be very low or on the contrary very high for several years in a row. Moreover, the harmfulness may change within a few years as resistances appear and more generally control practices are bypassed [3]. Classical estimates of harmfulness based on comparisons between treated on non-treated plots [4, 5] give us estimates of maximum losses induced by bioagressors. However, they do not inform us on what is the actual deficit of production due to bioagressors in the usual control practices. In addition, co-occurenceof pests and diseases make difficult to tease apart the effects of different bioagressors and other growing conditions such as weather and soil. Small scale studies then usually focus on the effect of one or a few pests or diseases [6, 7, 4].

Physiological mechanism models have the advantage of estimating a yield loss induced by one or several bioagressors and relatively to specific control practices, soil and weather defined achievable yields [1] but those estimates face calibration and validation difficulties [8], [9]. *Voir si il y a des choses sur la faillite mondiale de la prédiction de la production de blé en France en 2016. voir notamment Comparing and combining process-based crop models and statistical models with some implications for climate change et articles connexes (citant étant cités)*

Epidemiological survey data available for field crops, associated with yield data from the French Ministry of Agriculture represent a good opportunity to assess the correlations between bioagressors and yields, and thus the deficit of production possibly due to bioagressors activities under the usual control practices and weather conditions. Such large spatial scales strongly reinforce the efficiency of statistical models of yields *ref: https://www.sciencedirect.com/science/article/pii/S0168192310001978*. Moreover they provide observations on a large collection of bioagressors potentially having an important impact on yields. Here we focused on analysis on wheat, winter oilseed rape (WOSR) and barley, the three main grain crops in France. On those crops, we studied jointly the 25 bioagressors (13 diseases or 12 insects) most observed in a subset of the epidemiological survey data covering approximatively two third of grain crops surfaces in metropolitan France between 2009 and 2016 [10]. Once estimated the bioagressors impact on yields over the studied years we also estimated from the annual official reports of the French plant epidemiological surveillance authorities whether the pests and diseases studied had reached sufficiently high abundances during the years studied to be considered of concern.

## 2. Material and Methods

### 2.1 Quantification of bioagressor pressure per field and year

Observation data for pests and diseases are observations from the French epidemiological surveillance network gathered in the Vigicultures^®^ database [10]. It includes field crop monitoring carried out by various actors coordinated within the epidemiological surveillance network since 2009 in France (excluding administrative regions of Brittany, Pays de Loire, Limousin and Alsace). These weekly observations of plots during the growing season cover all insects and diseases considered a threat during the given period. As these measures are weekly and per plot, comparing them with departement level yield data requires aggregation by department and year. In line with the frequent use of nuisance thresholds, we counted in each plot how many times bioagressor counts exceeded bioagressor specific thresholds in a year.

#### 2.1.1. Choice of bioagressor metrics and thresholds

Several metrics are used in epidemiological surveillance databases to quantify the presence or harmfulness of a bioaggressor. For consistency, we selected the most commonly used metric in the surveys to quantify the bioagressor pressure. Three severity thresholds were tested for each pest or disease:

- Median threshold: For each plot-year, the mean of the observations is computed, then the chosen threshold correspond to the median of all of these values.
- Low and High threshold: When entering epidemiological surveillance data, indicative severity classes are provided to observers who use them in a variable way, reporting either absolute abundance values or values corresponding to the lower limits of these classes. We use here as a ‘Low’ threshold the input value in the second class and as a ‘High’ threshold the input value in the last class.

The corresponding metrics and thresholds are documented in Table 1.

**Table 1.** Summary of the metrics used to compute bio-agressor pressure

| Bio-agressor | Crop | Metric | Median threshold | Low threshold |
| --- | --- | --- | --- | --- |
| Gall midge | Winter wheat | Number of galls per trap | 0.67 | 5 |
| Slug | Winter wheat | Percentage of young plants damaged | 0 | 10 |
| Fusarium wilt | Winter wheat | Percentage of stems damaged | 0 | 10 |
| Silver scurf | Winter wheat | Percentage of the F3 leaf with symptoms | 0 | 2 |
| Barley powdery mildew | Winter wheat | Percentage of the F3 leaf with symptoms | 0 | 2 |
| Take-all | Winter wheat | Percentage of roots with necrosis | 0 | 10 |
| Bird cherry-oat aphid | Winter wheat | Percentage of plants damaged | 0 | 10 |
| Aphid | Winter wheat | Percentage of ears damaged | 0 | 10 |
| Eyespot | Winter wheat | Percentage of stems damaged | 0 | 10 |
| Brown rust | Winter wheat | Percentage of the F3 leaf with symptoms | 0 | 2 |
| Yellow rust | Winter wheat | Percentage of the F3 leaf with symptoms | 0 | 2 |
| Septoria leaf blotch | Winter wheat | Percentage of the F3 leaf with symptoms | 2.3 | 2 |
| Rape weevil | Rapeseed | Number of insects per trap | 0.57 | 1 |
| Cabbage seed weevil | Rapeseed | Number of insects per plant | 0.012 | 1 |
| Rape stem weevil | Rapeseed | Number of insects per trap | 1.6 | 1 |
| Rape flea beetle | Rapeseed | Number of insects per buried trap | 1.5 | 1 |
| White mold | Rapeseed | Percentage of flowers damaged | 45 | 30 |
| Pollen beetle | Rapeseed | Percentage of plants with insects | 20 | 25 |
| Cabbage aphid | Rapeseed | Number of aphids colonies per m<sup>2</sup> | 0 | 0 |
| Blackleg disease | Rapeseed | Percentage of necrosed area on the collar | 0 | 10 |
| Green peach aphid | Rapeseed | Percentage of plants with insects | 0 | 1 |
| Silver scurf | Winter barley | Percentage of the F3 leaf with symptoms | 1.5 | 2 |
| Barley scald | Winter barley | Percentage of the F3 leaf with symptoms | 1.16 | 2 |

*Utiliser multicolumn package sur les tables pour pouvoir aller à la ligne dans une cellule voir No Rapeseed, everything everywhere is WOSR (Winter Oilseed Rape)*

All data processing was done with the R software.

#### 2.1.2. Calculation of the annual bioagressor pressure at departmental scale

We tested two ways of aggregating at departmental level the weekly measures at plot level. The first is simply the rate of observations exceeding the chosen threshold in the department. However, this rate can be of low interest in some departments where observations are few in numbers. Indeed, in this case the pressure value is rapidly saturated either to 0 or 1. To limit this saturation, we also estimated the pressure as the result of a mixed binomial model of the rate of observations with a random effect on the interaction between the department and the year of observation (R package “glmer”).

### 2.2. Yield modelling

The annual yields at the departmental level for the different crops studied come from the Agreste database of the French Ministry of Agriculture and Food available online *ref*.

Large variations in yield for climatic reasons and regardless of the presence of pests and diseases could significantly blur the relationship between bioagressors and yield. For this purpose, estimates of achievable yield are commonly used, but their choice is still subject to discussion ([1]). Conversely, insects and plant diseases are strongly dependant on weather conditions, so incorporating weather factors in the regression could hide the direct impact of bioagressors. Rather than choosing one imperfect estimation, we test correlations between yield and bioagressors with or without proxies of achievable yields i.e, variables that are not necessarily significant in themselves but are strongely correlated to the achievable yield, which is neither observable nor directly measurable.

We test two conceptually different proxies of achievable yields based either on weather and soil information or on other crops yields. Estimating the variance explained by those proxies summarizing whether environmental conditions are favourable also allows to estimate the share of the departemental yield variance explained by environmental factors beyond bioagressors.

#### 2.2.1. Weather and soil based achievable yield

In a rather classic approach we first tried to characterize the achievable yield from weather and soil data. We used meteorological data of the French CNRM (Centre National de Recherches Météorologiques) from the SAFRAN objective analysis module [11]. Initially daily and per mesh, the different measured parameters were averaged per month and per department. We used the average daily precipitation (mm), the average daily evapo-transpiration (ETP, mm), the average daily radiation, the average daily maximum and minimum temperatures (^¤^C). In addition, we used GisSOL’s estimate of the useful water reserve per soil type [12]. The values, in mm, by soil type were averaged by department. To estimate achievable yields from these soil and meteorological data, we first performed a GAM-LASSO regression [?] on these monthly data in order to select the most relevant variables in the yield modeling by allowing a non-linear relationship between climate variables and yield:

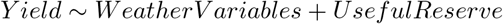

. We then used as achievable yield the predicted 9th decile of a quantile gam regression performed from the variables selected with the R package “qgam” *Add ref.*

##### 2.2.2. Use of other crops as proxies of achievable yield

We observed a significant correlation between the yields of the 3 winter crop species (oilseed rape, soft wheat and barley), commonly sown within 2 months in the fall and reaped within one month in the summer, exact dates varying by region and year. The departmental yield of barley alone explains more than 90% of the variations in wheat yield between 2009 and 2016, and about 60% of that of oilseed rape *add reference to graphs of correlation here* despite little to no overlap between the communities of bioagressors studied (only *oïdium in latin* is potentially common to wheat and barley.

Using an other crop yield as a covariate (i.e.: a proxy of achievable yield) could then allow to decorrelate yield residuals and generally favorable weather conditions allowing to explain those residuals by the presence of bioagressors.

##### 2.2.3. LASSO modeling of departmental yields

The yields were modeled as a linear regression between yield and bioagressors pressures, adding or not as covariates the proxies of achievable yield presented above. We then present the results in their diversity allowing to visualize their consistency and potential tendencies. To identify reliable correlations, we used a LASSO regression *cite original paper not the package paper* with selection of the regularization level (also performing variable selection) by cross validation with the R package “glmnet”[**?**]. Some bioagressors might be significantly positively correlated with the yields as they tend to be positively correlated with favorable conditions. As such correlations are likely spurious and could favor the emergence of other spurious correlations in the model we made use of the possibility in LASSO regressions to force the sign of selected correlations and present the results with or without this forcing.

Finally, as the ability to adequately control the bioagressors could vary with the region, we tested the the interaction between large agro-climatic regions (**Agro-climatic map**) and pest pressures has been taken into account for some models to highlight the presence of regional disparities on the effect of pests on yield.

#### 2.3. Analysis of official annual reports on bioagressors

Our estimates of bioagressors impact on yield are highly dependent on the presence of the bioagressors the studied years. To estimate if the studied years were representative of the known range of abundances for those bioagressors studied annual reports by the french regional Chambers of Agriculture. Such annual reports summarize the season newsletters published during the crop season. They give a detailed qualitative description of the intensity of bioagressors attack during the crop season, and often an estimation of corresponding maximum yield losses. Some annual reports are available online and thus give us very good references for our pressure calculations. We found four annual reports for the region “Ile-de-France” over the 2014-2017 period and three for the Picardie region (2014,2016,2017).

To convert the qualitative estimate given in annual reports into a quantitative assessment of the severity of the attack in the known range for each bioagressor we use a classic elicitation tool: the roulette (**Johnson et al., 2010 à citer**). This tool is integrated in a questionnaire that allows to document the arguments justifying the estimation by the roulette and to compare the different estimations, hear for the different years and regions (https://ecosys.versailles-grignon.inra.fr/SpatialAgronomy/compel).

With this questionnaire, we gave for each year a rate of intensity on a 1 to 10 scale for each bioagressor mentioned. These notations were then rescaled notations to compare the “calculated” pressure and the “reviewed” pressure.

## 3. Results

*Ajouter au moins dans les annexes les résultats utilisés pour les modèles qgam du rendement potentiel (1 par culture)*

The two-to-one correlations between bioagressors are low. On wheat, fusarium head blight and foot rot are the most correlated with a coefficient close to 0.4. On oilseed rape, only the two flea beetles show a significant correlation, with a coefficient close to 0.5. The two diseases of barley are not correlated. Consistently, the PCA does not reveal components that adequately describe all pressure variability: the main component explains only 18% of the variability of the 12 wheat bioagressors, 17% of the variability of the 9 oilseed rape bioagressors and 55% of the variability of the 2 barley bioagressors.

### 3.1. Comparison between “reviewed” and “calculated” pressures

The results (Figure 1) only concern “Ile-de-France” and “Picardie” regions over the 2014-2017 period. The link is shown between the reviewed pressure and the pressures calculated from epidemiological surveillance data. The pressure presented here were calculated with a mixt binomial model with the median threshold. Some bioagressors studied in the rest of the paper are not shown here as they were not mentioned in the season review.

**Figure 1.**
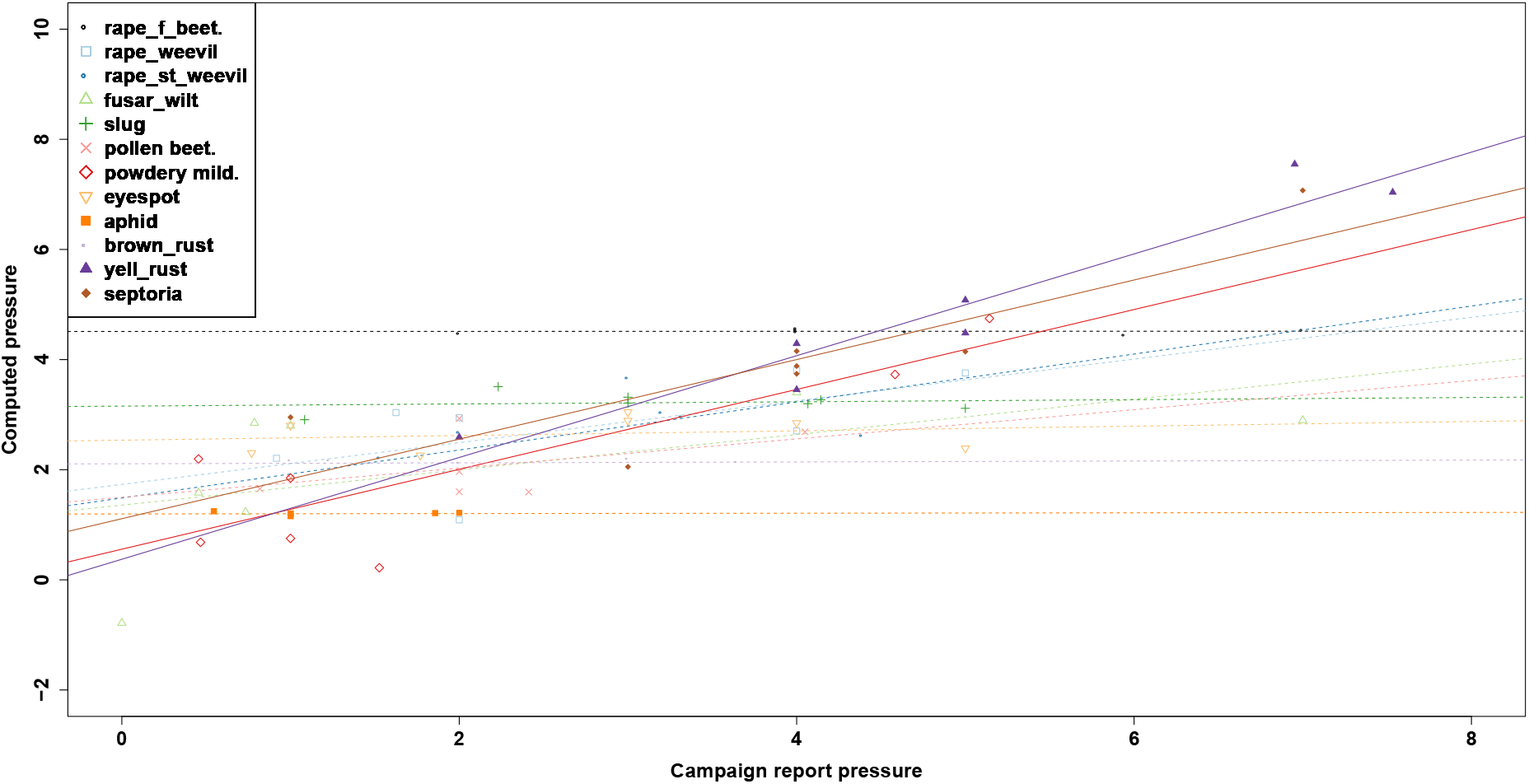
Corrlation between “calculated” pressures and “reviewed” pressures in Île de France and Picardie regions over the 2014-2017 period.

A positive and significant correlation between calculated and reviewed pressure were found for only three bioagressors (Figure 1): yellow rust, septoria tritici and powdery mildew. Variatons of calculated pressures seem to be smaller than those of reviewed pressures. However, when we rescale the calculated pressures on the “reviewed pressure” scale (Figure 2), we see that overall the calculated pressure cover the whole reviewed pressure scale. Our method to compute bioagressor pressure is *a priori* well suited to account for the variations of bioagressor pressures over the 2009-2016 period.

**Figure 2.**
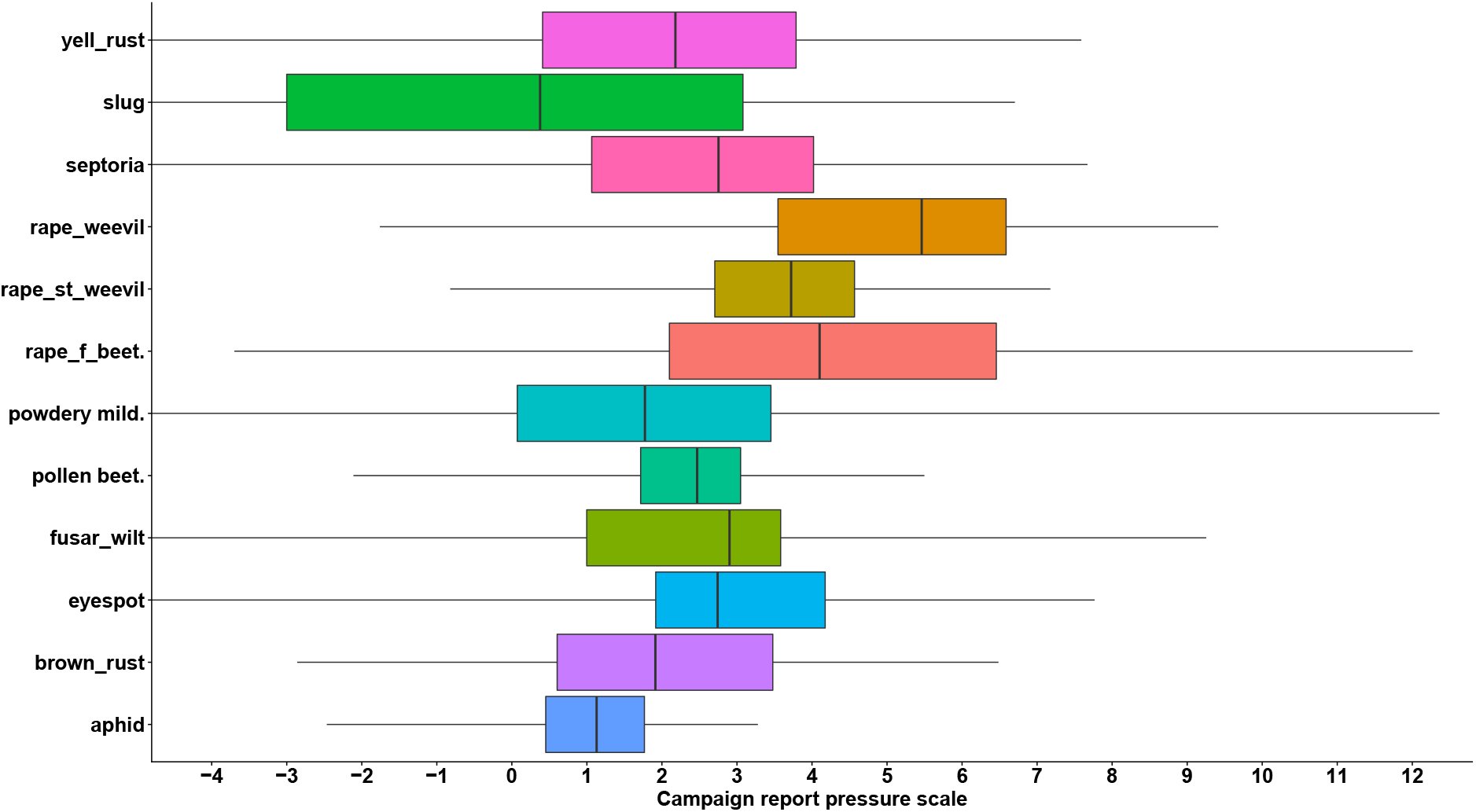
Boxplot comparing the scales of “calculated” and “reviewed” pressures for each bioagressor.

### 3.2. Results of yield modelling

#### 3.2.1. Winter wheat

On all models (La-___) and (Lb-___), only septoria and ear aphid show a negative, significant and recurrent correlation with winter wheat yield, with associated average losses of about 2 qx/ha (Figure 3). Other bioagressors, such as take-all, powdery mildew and yellow rust, are also present in some cases with a positive correlation in models whose bioagressors estimates are not forced to be negative or zero (La____), which can probably be explained by the presence of residual confounding effects related to meteorology.

**Figure 3.**
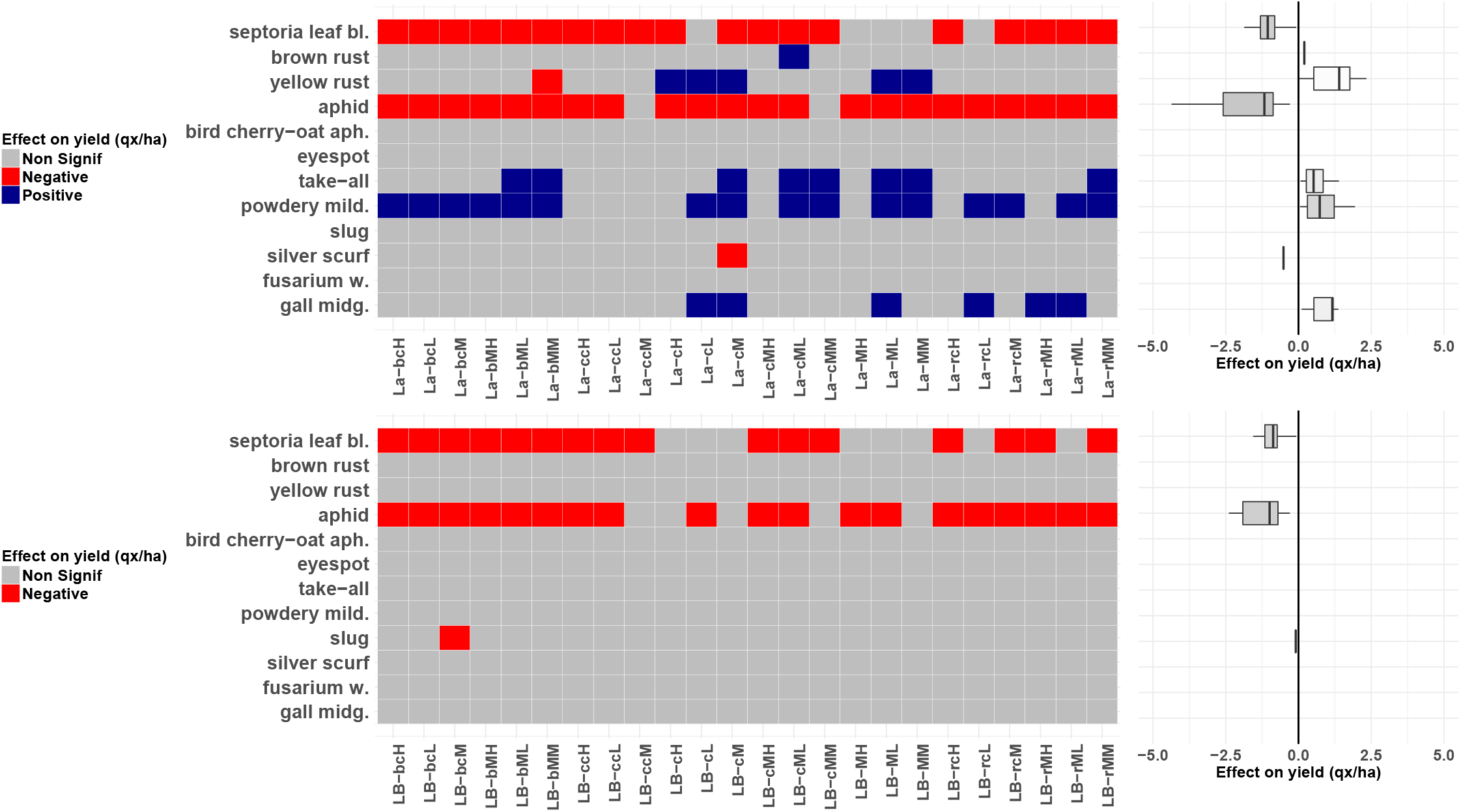
Synthesis matrix of the modelization results obtained on winter wheat. Pests are on the ordinate. Models are on the abscissa. The cells turn red when the pest pressure correlates significantly with the yield, otherwise gray. The box-plot graph represents the average losses associated in each model with pests. Model name: LB– 123: 1: Potential yield;, b: Barley, c: climatic, “”:none, r:oilseed rape. 2: Pression type; c: classical, M: mixed. 3: Threshold; L: Low, M: Median; H: High.

The predictive capacity of the models does not seem to be unreasonably affected by forcing bioaggressors to have a negative or no impact on yield (Figure 4). The total share of variation explained by the “LB” models is indeed very close to the equivalent “La” models. On the other hand, it depends largely on the proxy of the achievable yield used. It is between 80 and 85% with the “climate” proxy (models (La|LB)-c__), about 90% with the “barley” proxy (models (La|LB)-b__), between 50% and 60% with the “rape” proxy (models (La|LB)-r__) and about 5% without proxy (models (La|LB)-__). The percentage of yield variance explained by pests and diseases is higher in unbounded LASSO models (La), where it varies between 10% and 30%. It is generally higher when the threshold used is the low threshold. This percentage is more stable in bounded Lasso (LB) models, where it varies between 5 and 10%.

#### 3.2.2. Winter oilseed rape

For oilseed rape, no disease has a significant correlation with yield, regardless of the model (Figure 5). However, several insects emerge with a significant impact, in particular rape and cabbage seed weevils with average losses of about 1q/ha. This result confirms that the current losses due to oilseed rape diseases are limited compared to the losses caused by pests. Once again, we can see that the introduction of an achievable yield proxy is very important for the predictive capacity of our models (Figure 6). It alone explains about 40% of the variation in oilseed rape yield, a fairly stable percentage regardless of the proxy. However, this has little influence on the proportion of the variation explained by pests and diseases, which remains between 15% and 25%, except for a few models where no bioagressors are selected.

**Figure 4.**
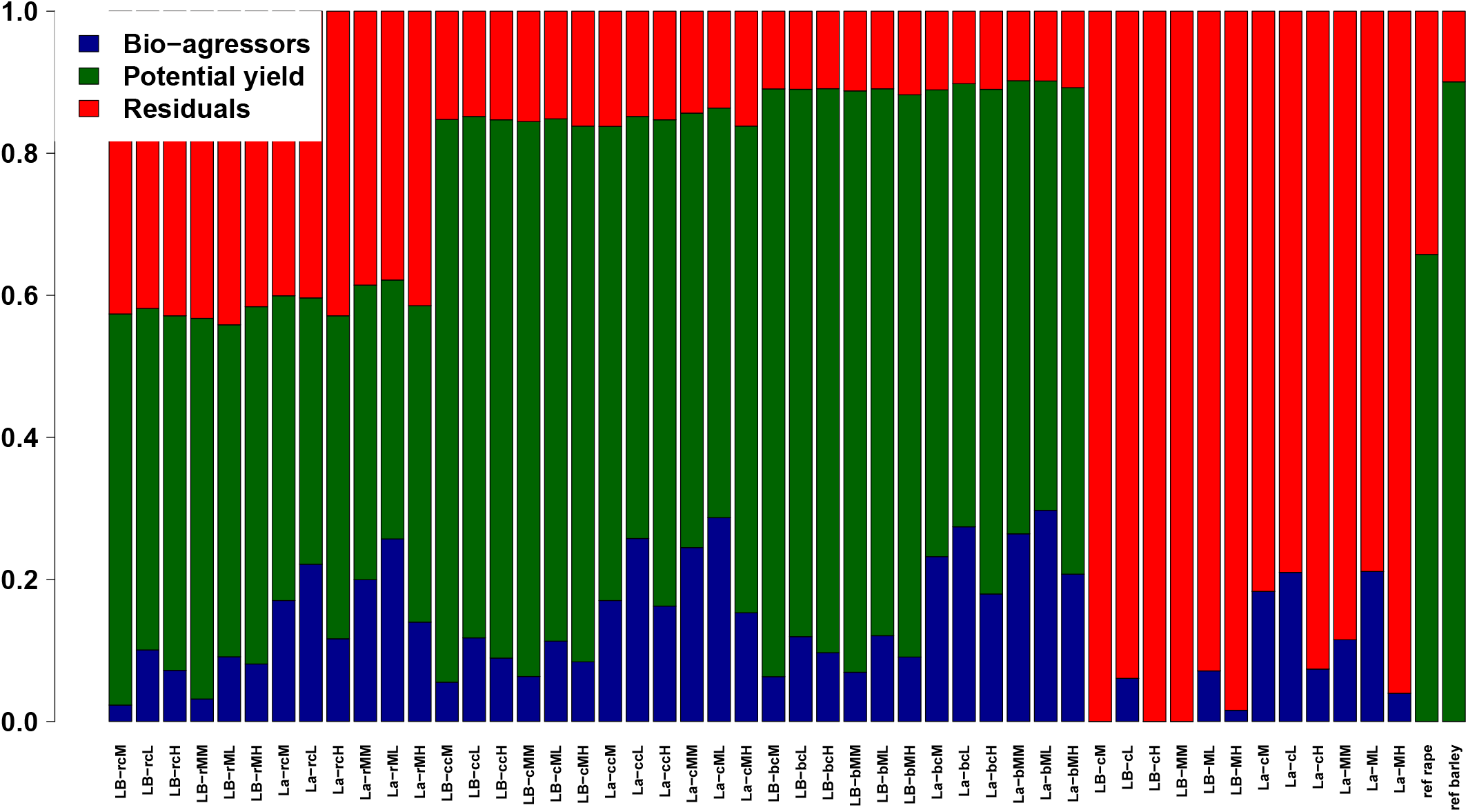
Share of the variance explained by each parameter in each model (winter wheat).

**Figure 5.**
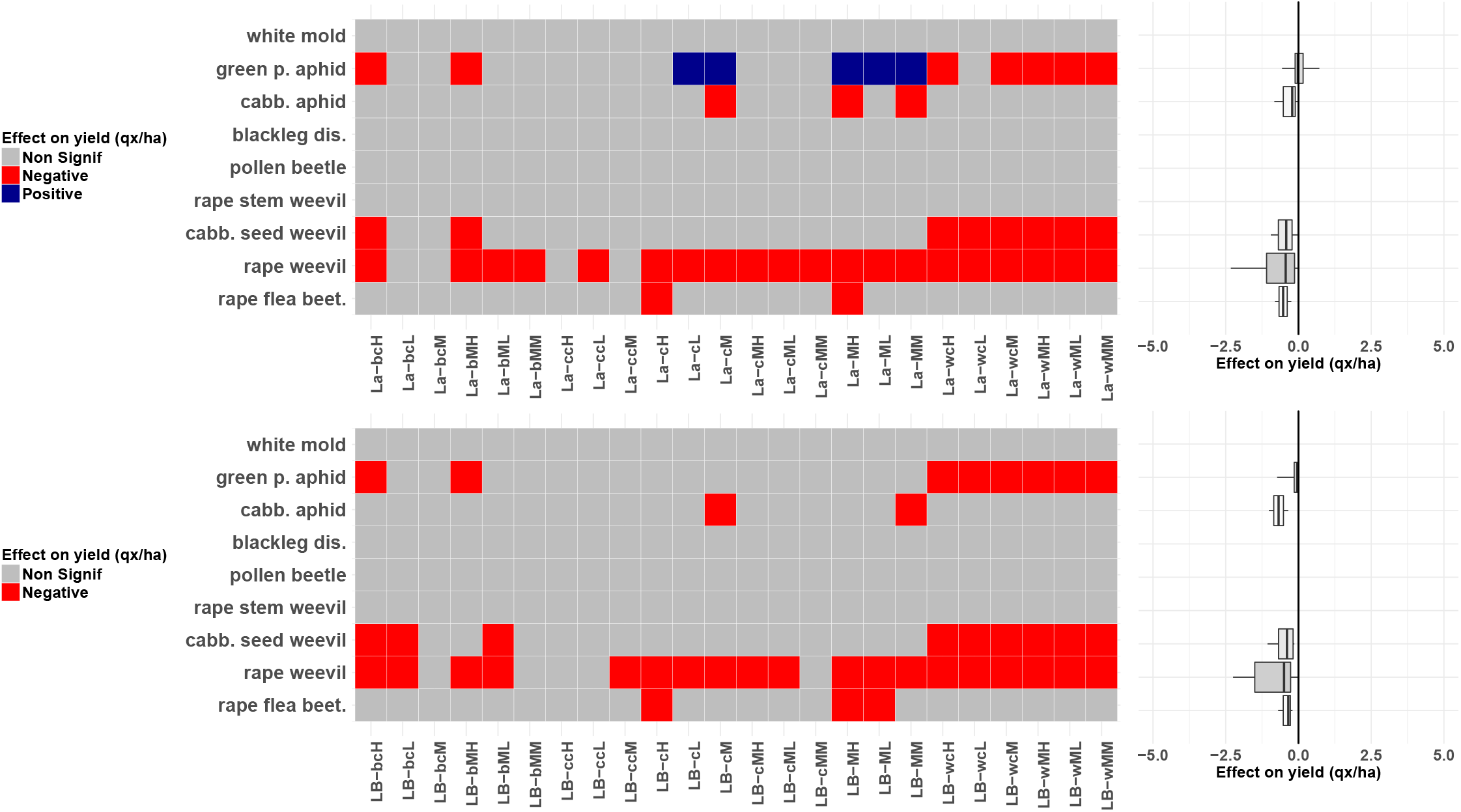
Synthesis matrix of the modelization results obtained on rape. Pests are on the ordinate. Models are on the abscissa. The cells turn red when the pest pressure correlates significantly with the yield, otherwise gray. The box-plot graph represents the average losses associated in each model with pests. Model name: LB– 123: 1: Potential yield;, b: Barley, c: climatic, ““:none, w:Wheat. 2: Pression type; c: classical, M: mixed. 3: Threshold; L: Low, M: Median; H: High.

**Figure 6.**
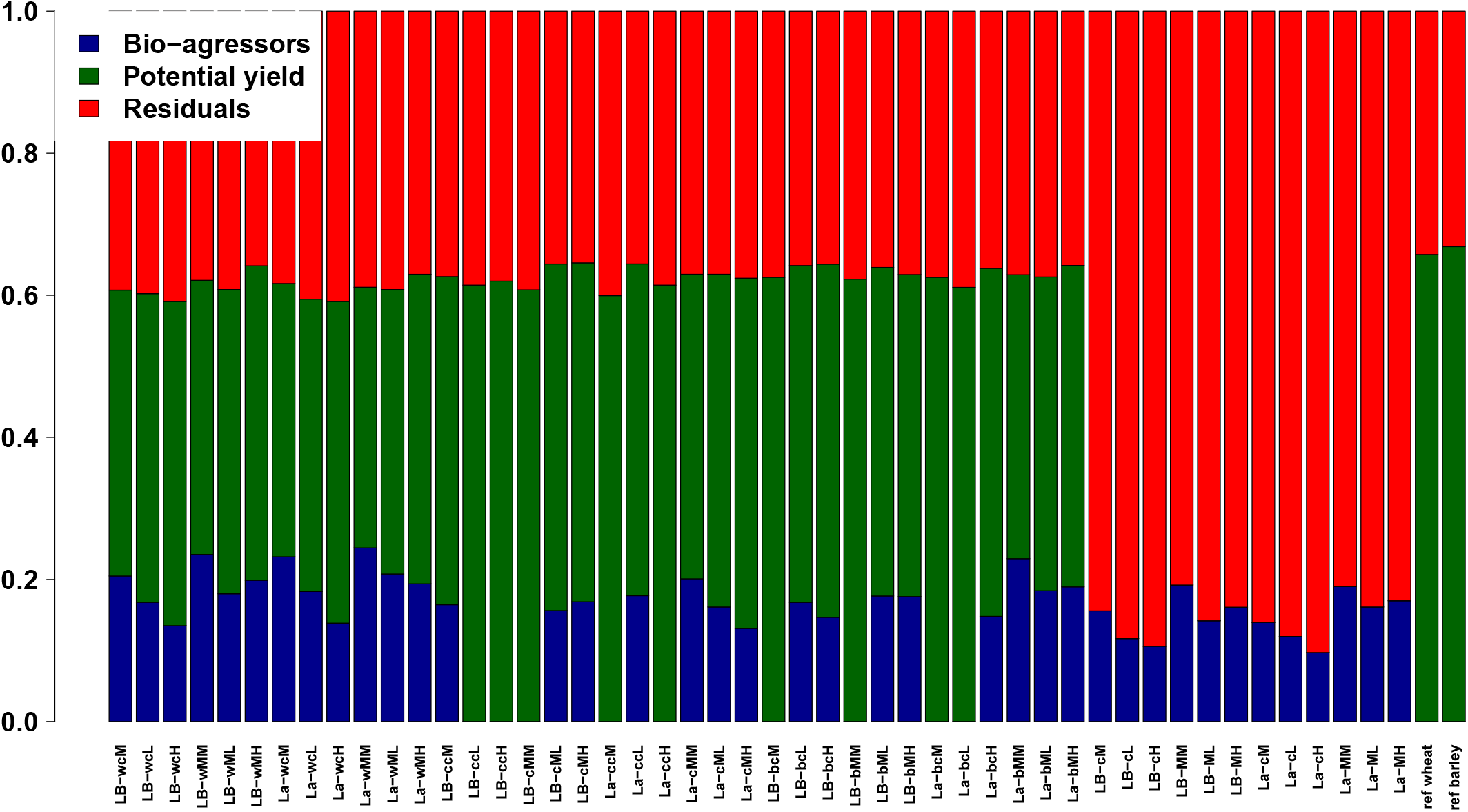
Share of the variance explained by each parameter in each model (winter oilseed rape).

#### 3.2.3. Winter barley

Of the two diseases studied on winter barley, none were correlated with yield. Rhynchosporiosis and helminthosporiosis appear to be well controlled.

## 4. Discussion

Our method enables to report yield losses at the department level caused by bioagressors. Septoria and ear aphids on wheat are responsible for average yield losses about 2 qtx/ha in France over the 2009-2016 period, corresponding to 5 to 10% of departmental yield variations of winter wheat. Those two bioagressors are also significant at the regional scale, particularly in western regions (Figure XXX).

Fusarium wilt also causes significant losses in the south-west and yellow rust minor losses in the center-east (Figure XXX). Losses due to oilseed rape and barley diseases are not significant neither on a national scale or a regional scale. On oilseed rape, only pests have a significant impact, which was expected since diseases of oilseed rape were not a major issue in recent years.

Other models predictive of the impact of pests and diseases have been developed, such as WHEAT-PEST ([8]), or CERES-Rice ([13]), respectively on soft wheat and rice. These are agrophysiological mechanistic models. The achievable yield of the crop is first estimated using a growth equation, with a daily time step. It depends on meteorological variables (daily temperature, radiation, etc.), soil and agricultural practices (excluding treatment). These models determine the injury profile from the agricultural and climate context.

Unlike these models, the work carried out is based exclusively on the statistical analysis of large datasets, limiting as much as possible the assumptions about the functioning of the crop-bioagressor system. In particular, we show that the yield of another crop with a similar vegetation cycle can be an excellent proxy for yield and acts as a biological “model” simply replacing computer models. Slightly more complex, our statistical yield modeling based on meteorological and soil data also provides excellent yield approximations. While each of the achievable yield proxies we use is questionable individually, the broad concordance of the results obtained despite the diversity of these proxies suggests that our results are highly robust to the exact formulation of the achievable yield.

In addition, mechanistic yield models are commonly used to estimate maximum losses associated with the presence of a pest infestation. Taking treatments into account is difficult because the effectiveness of the treatments requires additional parameter setting of the model, which is particularly difficult because the effectiveness is subject to variation over time. More generally, the difficulty of setting mechanistic models for the whole diversity of situations present in France contrasts with the simplicity of the statistical method proposed here, which should make it possible to continuously assess whether a disease or an insect is or becomes harmful to crops. We estimated the production deficit related to Septoria in treated systems at 2 qx/ha, which may seem low compared to the 17qx/ha advanced by the technical institutes in untreated plots(http://www.fiches.arvalis-infos.fr/fiche_accident/fiches_accidents.php?mode=fa&type_cul=1&type_acc=4&id_acc=46). Similarly, other diseases did not emerge as having a significant impact on yields. However, such differences were expected since we are interested in actual losses considering the treatments usually performed and not in potential losses without treatment. It is also important to note that our results are an average over a period of 8 years and at the departmental level while bioagressor pressures vary significantly between plots and especially from one year to the next. Nevertheless, the comparison of our “calculated” pressure with the “reviewed” pressure remains encouraging, since the two scales of variation correspond relatively well (Figure 2). A priori, the pressure variations at the departmental level over the study period are well taken into account, which is essential to determine the associated production deficits.

*Comment achievable yields main parameters in the light of Tamara’s paper: https://www.sciencedirect.com/science/arti*

Our method makes it possible to overcome the differences in metrics between bioaggressors and to introduce them together into the models, which is important because all can potentially have an impact on yields. On the other hand, the fact that a bioaggressor pressure is significantly correlated with yield losses could depend on the metric used to quantify its abundance. In particular, if other metrics are better correlated to plot damage than those most commonly used in the epidemiological surveillance network, they could also be significantly correlated to variations in departmental yields. For example, the abundance of pollen beetles in a given year is not directly correlated to the damage caused, but will also depend on the correspondence between the insect’s flight period and the crop’s sensitivity period.

Overall, the achievable yield proxies used explain a larger share of the yield variation of a crop than pests and diseases, suggesting that yield variations on wheat, oilseed rape and barley over the 2009-2016 period are more related to weather than to the attack of the bioagressors themselves.

## 5. Conclusion

The majority of bioaggressors in winter barley, oilseed rape and winter wheat were not significant determinants of yield at the departmental level between 2009 and 2016. On the other hand, a control deficit seems to be observed on wheat with septoria and ear aphid, about 2 qx/ha each, and on oilseed rape with rape and cabbage seed weevils. In a rather original way, the calculation method used made it possible to evaluate the residual impact of pests and diseases in a chemical control situation. It could be used for national continuous monitoring of resistance breaking and more generally deficiencies in the management of field crop bioagressors.

**Table 2.** Links to find the regional crop season reports for Ile de France and Picardie over the 2014-2017 period

| Region | year | Link |
| --- | --- | --- |
| Ile -de-France | 2014 | http://driaaf.ile-de-france.agriculture.gouv.fr/IMG/pdf/BILAN_SANITAIRE_grandes |
| Ile -de-France | 2015 | http://driaaf.ile-de-france.agriculture.gouv.fr/IMG/pdf/BILAN_SANITAIRE_2015_-_ |
| Ile -de-France | 2016 | http://driaaf.ile-de-france.agriculture.gouv.fr/IMG/pdf/2016_GC_BILAN_SANITAIRE |
| Ile -de-France | 2017 | http://driaaf.ile-de-france.agriculture.gouv.fr/IMG/pdf/GC_2017_BILAN_SANITAIRE |
| Picardie | 2014 | http://blog-ecophytohautsdefrance.fr/wp-content/uploads/2015/12/Bilan-VF-sanita |
| Picardie | 2015 | http://www.hautsdefrance.chambres-agriculture.fr/fileadmin/user_upload/Hauts-de |
| Picardie | 2017 | https://hautsdefrance.chambres-agriculture.fr/fileadmin/user_upload/Hauts-de-Fr |

## 6. Appendices

## References

1. S. Savary, A. Ficke, J.-N. Aubertot, C. Hollier, Crop losses due to diseases and their implications for global food production losses and food security 4 (4) (2012) 519–537. doi:10.1007/s12571-012-0200-5. URL http://link.springer.com/10.1007/s12571-012-0200-5

2. D. C., D. O., M. A., C. M., Falgas., G. C.A., G. J.D., Laudinot., S. A., W. E., Conseil agronomique et réduction des pesticides: quelles ressources pour affronter ce nouveau challenge professionnel ? 34 (2014) 367–378.

3. A. R. Jutsum, S. P. Heaney, B. M. Perrin, P. J. Wege, Pesticide resistance: assessment of risk and the development and implementation of effective management strategies† 54 (4) (1998) 435–446. doi:10.1002/(SICI)1096-9063(199812)54:4<435::AID-PS844>3.0.CO;2-K. URL http://doi.wiley.com/10.1002/%28SICI%291096-9063%28199812%2954%3A4%3C435%3A%3AAID-PS844%3E3.0.CO%3B2-K

4. S. Savary, P. S. Teng, L. Willocquet, F. W. Nutter, Quantification and modeling of crop losses: Areview of purposes 44 (1) (2006) 89–112. doi:10.1146/annurev.phyto.44.070505.143342. URL http://www.annualreviews.org/doi/10.1146/annurev.phyto.44.070505.143342

5. C. Chevalier-Gérard, J. Denis, J. Meynard, Perte de rendement due aux maladies cryptogamiques sur blé tendre d’hiver. construction et validation d’un modèle de l’effet du système de culture 14 (5) (1994) 305–318. doi:10.1051/agro:19940504. URL http://www.agronomy-journal.org/10.1051/agro:19940504

6. W. Padwick G., Losses causes by plant diseases in the colonies.

7. M. R. McRoberts N., Franke A.C., Survey of phalaris minor in the indian rice-wheat system. 39.

8. L. Willocquet, F. A. Elazegui, N. Castilla, L. Fernandez, K. S. Fischer, S. Peng, P. S. Teng, R. K. Srivastava, H. M. Singh, D. Zhu, S. Savary, Research priorities for rice pest management in tropical asia: A simulation analysis of yield losses and management efficiencies 94 (7) 672–682. doi:10.1094/PHYTO.2004.94.7.672. URL http://apsjournals.apsnet.org/doi/10.1094/PHYTO.2004.94.7.672

9. J. Timsina, E. Humphreys, Performance of CERES-rice and CERES-wheat models in rice-wheat systems: A review 90 (1) 5–31. doi:10.1016/j.agsy.2005.11.007. URL http://linkinghub.elsevier.com/retrieve/pii/S0308521X05002623

10. F. N. N. of Plant Epidemiological Surveillance, Epidemiological suveillance data, aggregrated by department, http://www.vigicultures.fr.

11. N. C. o. M. R. CNRM, Weather data calculated from SAFRAN model, aggregated by department, https://www.umr-cnrm.fr/spip.php?article788.

12. I. GisSol, Water useful reserve data, aggregated by department, https://www.gissol.fr/.

13. H. Pinnschmidt, W. Batchelor, P. Teng, Simulation of multiple species pest damage in rice using CERES-rice 48 (2) 193–222. doi:10.1016/0308-521X(94)00012-G. URL http://linkinghub.elsevier.com/retrieve/pii/0308521X9400012G

